# Frequency-Aware Interpretable Deep Learning Framework for Alzheimer’s Disease Classification Using rs-fMRI

**DOI:** 10.1101/2025.09.18.677114

**Authors:** Yutong Gao, Robyn L. Miller, Vince D. Calhoun

**Affiliations:** Tri-Institutional Center for Translational Research in Neuroimaging and Data Science (TReNDS), Georgia State University, Georgia Institute of Technology, Emory University, Atlanta, GA, USA

**Keywords:** Deep Learning, Explainable AI, Frequency-Aware, Brain Dynamics, rs-fMRI

## Abstract

Gaining insight into the spectral and temporal alterations in brain connectivity associated with Alzheimer’s disease (AD) may offer pathways toward more informative biomarkers and a deeper understanding of disease mechanisms. We propose FINE (Frequency-aware Interpretable Neural Encoder), a novel deep learning model designed to capture multi-scale temporal and frequency-specific patterns in dynamic functional network connectivity (dFNC) derived from resting-state fMRI. FINE integrates multiple expert branches, including convolutional layers, learnable wavelet layers, transformers, and static encoders, enabling the joint modeling of temporal evolution and spectral content of brain networks in an end-to-end framework. Beyond classification, FINE supports frequency-wise interpretability by aligning gradient-based saliency maps with statistical group differences, revealing potential robust, biologically meaningful biomarkers of AD. Evaluated on the large OASIS-3 dataset (856 subjects), FINE achieves AD classification performance (ROC-AUC 0.769) and provides insights into frequency-specific connectivity disruptions, particularly within subcortical, sensorimotor, and cerebellar networks. Our results demonstrate that incorporating frequency-aware modeling and interpretable architectures can advance both disease classification and underlying functional disruption of AD-related brain dynamics.

## I. Introduction

The human brain functions as a richly dynamic and intricately organized system. Blood oxygenation-level dependent (BOLD) functional magnetic resonance imaging (fMRI) captures variations in blood flow, providing an indirect proxy for localized neuronal activity. Investigating brain dynamics using resting-state fMRI (rs-fMRI) has substantially enhanced our understanding of neural processes underlying aging, sex differences, and cognitive disorders.

Functional connectivity (FC) [1], which reflects correlations between intrinsic brain regions, captures the brain’s evolving network organization. Using group independent component analysis (GICA), the brain is decomposed into a set of independent component networks (ICNs), each representing a distinct functional domain with its own temporal profile. Numerous methods have been proposed to estimate time-resolved functional network connectivity (FNC) from ICNs’ time courses, where dynamic correlations reflect the evolving interactions among brain networks. Most of these methods operate at a fixed temporal scale. For instance, static FNC assumes constant connectivity throughout the acquisition period [2], while sliding-window dynamic FNC applies a fixedsize temporal window to capture time-varying patterns [3]. Alternatively, windowless wavelet-based approaches use complex Morlet wavelets to estimate instantaneous changes in connectivity [4]. However, both sliding-window and wavelet-based methods typically rely on predefined temporal resolutions, which may limit their ability to fully capture the brain’s multi-scale dynamics. To address this limitation, some studies have introduced frequency-resolved approaches that explicitly model connectivity across multiple spectral bands. For example, [5] proposed a filter bank-based framework to decompose FNC into band-limited components, while dynamic amplitude of low-frequency fluctuation (ALFF) analyses focus on specific frequency bands to reveal group-level abnormalities, such as in schizophrenia [6]. These approaches aim to move beyond single-scale analysis by uncovering frequency-dependent patterns of brain network interactions.

Despite these advances, sliding-window Pearson correlation remains the most widely used method due to its simplicity and the board application of Pearson correlation itself [7]. Typical window lengths range from 30 seconds to 1 minute in rs-fMRI studies [1], [8]. Smaller windows allow detection of faster changes but act as a high-pass filter, attenuating slower fluctuations. Conversely, larger windows better preserve low-frequency information but may obscure meaningful high-frequency dynamics. For example, [9] and multiple ALFF studies emphasize the importance of low-frequency content in brain activity, while [10] argue that window sizes over 100 seconds may be needed to capture fluctuations around 0.01 Hz. Consequently, identifying reliable frequency-specific connectivity patterns using fixed-window correlation remains challenging and motivates the need for new models capable of learning from the full spectral and temporal richness of dFNC data.

Deep learning methods have recently made notable advances in brain imaging analysis. Architectures include convolutional neural network (CNN)-based models for schizophrenia classification using dFNC [11], LSTM-based forecasting from ICNs to predict future activity [12], and graph-based approaches for early-stage AD detection from spatial and temporal dFNC correlations [13]. Additionally, Brain Language Model (BrainLM), a transformer-based foundation model, has been introduced for modeling brain activity dynamics. Deep learning demonstrates strong capability in learning from high-dimensional and high-complexity transformed brain features. While spatial and temporal features have been extensively studied using deep learning, frequency-aware features, and their integration with spatial and temporal information remain underexplored.

Moreover, as deep learning models grow in complexity and predictive power, it becomes increasingly important to understand why and how they make their predictions, to uncover the inner workings of these models, and to gain insights into their outputs in data-driven systems, particularly in the context of disease-related brain imaging. In understanding brain mechanisms, where the insights from data and the tasks solved by models often remain opaque as model complexity increases, it becomes essential to provide concrete ways to interpret both the data and the model. Explainable AI (XAI) spans a broad spectrum of techniques, ranging from simple coefficient inspection in linear models and logical decision paths in decision trees to more advanced strategies involving explainable neural networks. These include posthoc approaches such as gradient-based and perturbation-based methods, as well as model-intrinsic approaches. For example, employed a gradient-based XAI method to identify potential biomarkers from dFNC features using a trained time-attention LSTM model. Non post-hoc approach such as, [16] utilized interpretable coefficient maps from GraphNet, applying structured regularization to highlight meaningful signal patterns across whole-brain fMRI data.

To address the limited exploration of frequency-aware deep learning models and their interpretability in frequency-sensitive domains, we propose FINE (Frequency-aware Interpretable Neural Encoder), a deep learning model built on a multi-expert architecture. FINE consists of parallel frequency-aware, structural branches, enabling the model to jointly learn dFNC patterns and their spectral characteristics in an end-to-end manner. The frequency-aware experts comprise modules such as CNNs with varying kernel sizes or Morlet wavelet transforms, which specialize in extracting frequency-specific features. In contrast, the structural experts, which may include transformers or linear layers, capture temporal dynamics and spatial connections where arises naturally from the correlation patterns within functional connectivity. Unlike traditional approaches that require prior frequency decomposition, FINE directly processes dFNC as input, allowing the model to learn both the temporal evolution and the underlying spectral components of brain connectivity. This design not only facilitates more effective feature learning but also enhances interpretability, as FINE supports post hoc saliency analysis and feature attribution to uncover frequency-specific biomarkers relevant to AD recognition.

### Contributions

In summary, our key contributions are three-fold: (1) we propose **FINE**, a novel multi-branch architecture for joint temporal and frequency-specific modeling of dFNC; (2) we introduce an interpretable framework that aligns deep model saliency with classical group-level statistics, providing frequency-wise biomarkers of AD; (3) we demonstrate empirically that frequency-aware modeling substantially improves AD classification and yields biologically consistent, frequency-specific insights into altered brain dynamics. We believe that FINE provides a flexible and extensible foundation for XAI in frequency-aware neuroimaging.

## II. Material and Methodology

In this section, we present *FINE (Frequency-aware Interpretable Neural Encoder)*, a deep learning model designed as a multi-expert architecture that incorporates parallel frequency-aware and structural branches. The frequency-aware modules (e.g., CNNs with varying kernel sizes or Morlet wavelet transforms) specialize in capturing spectral features, while the structural modules (e.g., transformers or linear layers) model temporal and spatial dependencies. The input to our model consists of dFNC matrices, computed using a sliding-window correlation approach across 53 ICNs derived from GICA via the Neuromark pipeline [3]. FINE aims to extract complementary representations from the dFNC sequences that are critical for identifying biomarkers in AD. The data processing pipeline is illustrated in Figure 1 and the model architecture of FINE is illustrated in Figure 2.

**Fig. 1:**
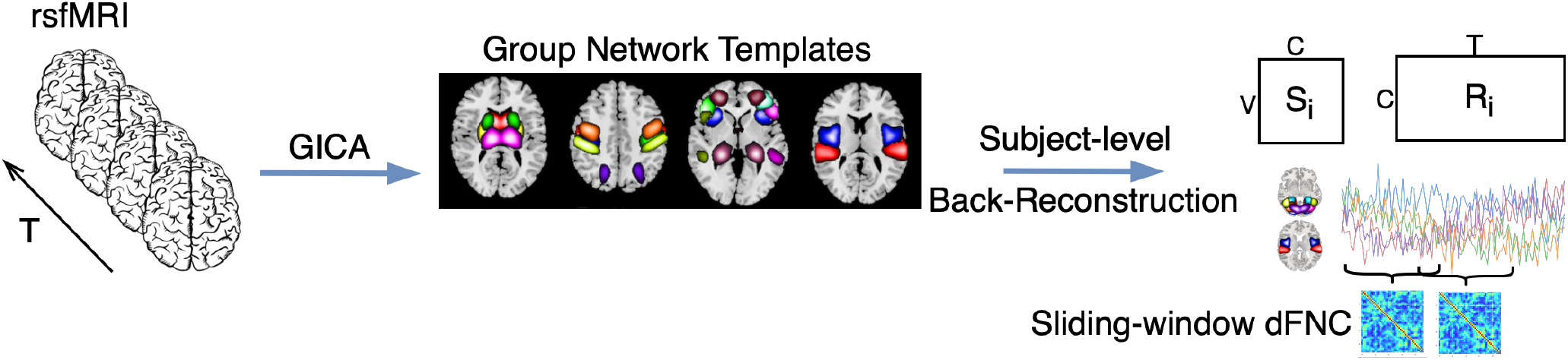
Schematic of the rs-fMRI data processing pipeline (figure modified from [1]). Preprocessed fMRI data are decomposed into 53 intrinsic connectivity networks (ICNs) using Neuromark [3], guided by replicated group-level templates from HCP and GSP datasets. Subject-specific spatial maps (*S*_*i*_) and time courses (*R*_*i*_) are derived via adaptive ICA. dFNC is computed using sliding-window Pearson correlation across ICN time series, resulting in a sequence of 139 correlation matrices per subject.

**Fig. 2:**
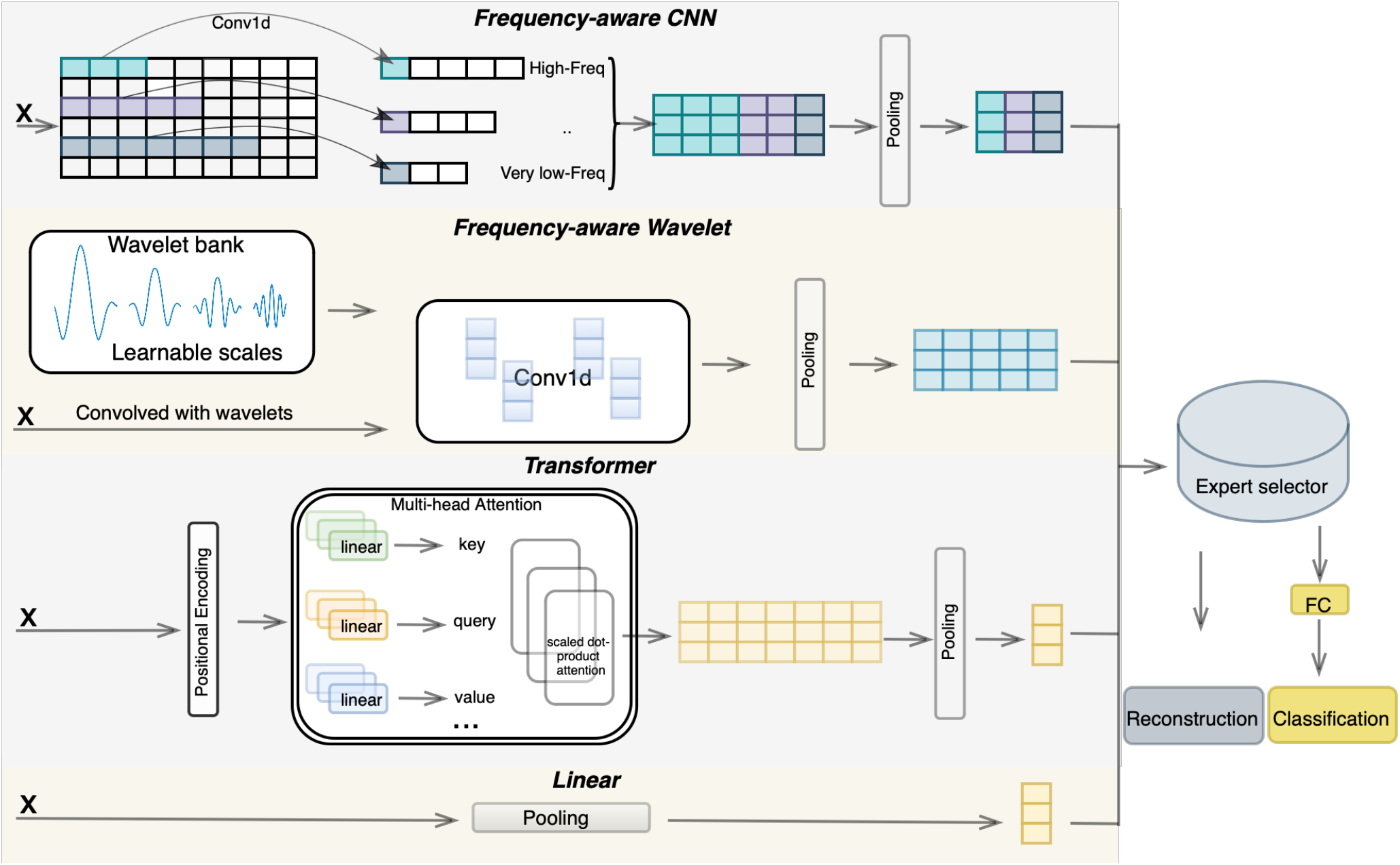
Overview of the proposed multi-expert architecture (**FINE: Frequency-aware Interpretable Neural Encoder**). The model consists of four parallel branches designed to capture complementary frequency and temporal dynamics from input dFNC sequences **X** ∈ ℝ^*T* ×*F*^. **Top:** The *frequency-aware CNN branch* uses multiple depthwise 1D convolutions with varying kernel sizes (e.g., 5 to 49) to approximate different frequency bands (from high to very-low), followed by adaptive pooling. **Second:** The *wavelet branch* performs convolution with learnable Morlet-like wavelet filters at multiple scales to extract spectral features, which are subsequently pooled. **Third:** The *Transformer branch* captures global temporal dependencies via multi-head self-attention. Positional encodings are added before feeding the sequence into the Transformer layers. **Bottom:** The *linear branch* applies global average pooling over time to extract static features (full-band FNC). The outputs from selected branches are concatenated via an *expert selector* module, allowing certain experts to be enabled or disabled during training or inference. The fused representation is used for two downstream tasks: input reconstruction and AD classification.

### A. Resting-state fMRI data

The data used in this study come from the Open Access Series of Imaging Studies (OASIS)-3 [17], a publicly available dataset collected from several ongoing studies conducted at the Washington University Knight Alzheimer Disease Research Center, with Institutional Review Board approval. The dataset includes samples across various stages of cognitive decline, comprising longitudinal rs-fMRI scans along with corresponding clinical and demographic information.

In this work, we focus on two groups: the cognitively normal (CN) group, defined as subjects with a Clinical Dementia Rating (CDR) score of 0, and the AD group, which includes subjects with CDR ≥1 as well as those with CDR = 0.5 who received a clinical diagnosis of AD dementia at the time of the scan [17]. A total of 856 unique subjects are included in our analysis, consisting of 654 CN subjects and 202 AD subjects. Given the longitudinal nature of the dataset, multiple scans may exist per subject, potentially across different disease stages. To avoid data leakage and ensure proper generalization, only one scan per subject is included in the study, with no subject appearing in more than one of the training, validation, or test sets. Demographic details of the included sample are summarized in Table 1. Given that AD is inherently age-related, the age distribution between CN and AD groups is naturally imbalanced in the dataset. In this study, our primary objective is to develop data-driven models for AD classification, without applying explicit age balancing or covariate adjustments. The age-related differences reflect real-world clinical distributions and are considered part of the disease phenotype as captured by the model.

**TABLE I:**
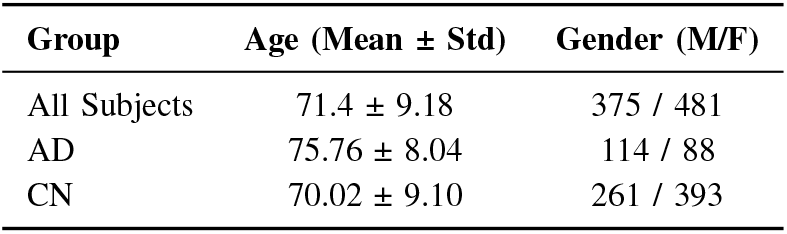
Demographic Summary of the Whole Dataset.

| Group | Age (Mean ± Std) | Gender (M/F) |
| --- | --- | --- |
| All Subjects | 71.4 ± 9.18 | 375 / 481 |
| AD | 75.76 ± 8.04 | 114 / 88 |
| CN | 70.02 ± 9.10 | 261 / 393 |

### B. Data Preprocessing

The rs-fMRI data were preprocessed using Statistical Parametric Mapping (SPM12) to correct for timing differences in slice acquisition. To correct for subject head motion, rigid body motion correction was first applied, followed by slicetiming correction. The fMRI volumes were then normalized to the standard Montreal Neurological Institute (MNI) space using an echo planar imaging (EPI) template and resampled to isotropic voxels with a resolution of 3 × 3 × 3 mm^3^. The resampled images were subsequently smoothed using a Gaussian kernel with a full width at half maximum (FWHM) of 5 mm. Figure 1 illustrates the schematic of applying Neuromark [3] for decomposing processed rs-fMRI data into ICNs. Group-level ICNs were obtained from two independent datasets: the Human Connectome Project (HCP) and the Genomics Superstruct Project (GSP). Highly replicated ICNs were identified by computing the correlation between their spatial maps across datasets. Subject-specific ICNs and their corresponding time courses (TCs) were then estimated through adaptive ICA, using these group-level templates as priors during the back-reconstruction process. A total of 53 ICNs were identified as highly reliable and functionally meaningful. These components were categorized into seven distinct functional domains, informed by anatomical location and established functional roles. The seven domains include the subcortical network (SC), auditory network (AU), sensorimotor network (SM), visual network (VI), cognitive control network (CC), default-mode network (DM), and cerebellar network (CB). Table 2 provides a detailed list of all 53 ICNs and their corresponding functional domain assignments.

**TABLE II:** Independent Component Networks (ICNs) Grouped by Functional Domain.

|  |  |
| --- | --- |
| **Subcortical network (SC)** | **Sensorimotor network (SM)** |
| Caudate (69) | Postcentral Gyrus (3) |
| Subthalamus (53) | Left Postcentral Gyrus (9) |
| Putamen (98) | Paracentral Lobule (2) |
| Caudate (99) | Right Postcentral Gyrus (11) |
| Thalamus (45) | Superior Parietal Lobule (27) |
| **Auditory network (AU)** | Paracentral Lobule (54) |
| Superior Temporal Gyrus (21) | Precentral Gyrus (66) |
| Middle Temporal Gyrus (56) | Superior Parietal Lobule (80) |
| **Visual network (VI)** | Postcentral Gyrus (72) |
| Calcarine Gyrus (16) | **Cognitive control network (CC)** |
| Middle Occipital Gyrus (5) | Inferior Parietal Lobule (68) |
| Middle Temporal Gyrus (62) | Insula (33) |
| Cuneus (15) | Superior Medial Frontal Gyrus (43) |
| Right Middle Occipital Gyrus (12) | Inferior Frontal Gyrus (70) |
| Fusiform Gyrus (93) | Right Inferior Frontal Gyrus (61) |
| Inferior Occipital Gyrus (20) | Middle Frontal Gyrus (55) |
| Lingual Gyrus (8) | Inferior Parietal Lobule (63) |
| Middle Temporal Gyrus (77) | Left Inferior Parietal Lobule (79) |
| **Default-mode network (DM)** | Supplementary Motor Area (84) |
| Precuneus (32) | Superior Frontal Gyrus (96) |
| Precuneus (40) | Middle Frontal Gyrus (88) |
| Anterior Cingulate Cortex (23) | Hippocampus (48) |
| Posterior Cingulate Cortex (71) | Left Inferior Parietal Lobule (81) |
| Anterior Cingulate Cortex (17) | Middle Cingulate Cortex (37) |
| Precuneus (51) | Inferior Frontal Gyrus (67) |
| Posterior Cingulate Cortex (94) | Middle Frontal Gyrus (38) |
| **Cerebellar network (CB)** | Hippocampus (83) |
| Cerebellum (13) |  |
| Cerebellum (18) |  |
| Cerebellum (4) |  |
| Cerebellum (7) |  |

#### 1) Functional Network Connectivity

The connectivity between the preprocessed time courses was calculated using Pearson correlation, resulting in a total of 1378 whole-brain correlations among 53 ICNs. The dynamic FNC was segmented using a tapered sliding window, generated by convolving a rectangular window (window size = 20, TR = 44s) with a Gaussian kernel (*σ* = 3), with a step size of 1 TR, producing 139 windows in total.

### C. Model Architecture

#### 1) Encoder - CNN Branch

The CNN encoder branch is aim to capture temporal patterns in the dFNC series at multiple scales. We employ a sequence of 1D convolutional layers with varying kernel sizes to extract frequency information, where smaller kernels capture high-frequency patterns and larger kernels capture low-frequency temporal dynamics. Specifically, the branch applies convolutions with kernel lengths *K* = 49, 27, 9, 5 in parallel. A larger kernel (e.g., 49) covers a longer temporal window, effectively capturing slow, long-duration fluctuations (*<* 0.01 Hz) in connectivity, whereas a smaller kernel (e.g., 5) focuses on rapid, short-term high-frequency changes. By using multiple convolutional layers with these kernel sizes, the network simultaneously learns multi-scale features. Each depthwise convolution uses separate filters on each of the 1378 connectivity channels independently, without mixing information across channels, thereby preserving the identity of each functional connectivity component. Non-linear activations ReLU are applied after each convolution.

To aggregate these feature representations, we concatenate **h**^Conv^ ∈ ℝ^*T* ×*C*^, where Conv includes very slow, slow, mid, and high-frequency feature maps produced by the depthwise CNN layers, *C* is the number of channels, and *T* is the sequence length. This representation summarizes the entire sequence’s features as learned by the CNN branch. Leveraging the multi-scale kernels, **h**^Conv^ captures frequency-specific characteristics of the dFNC sequence, including periodicities and quick bursts, and serves as an informative representation for subsequent decoding and classification.

#### 2) Encoder - Wavelet Branch

To extract frequency-specific representations from the input signal, we introduce a *Learnable Wavelet Layer*, which implements a Morlet-like wavelet transform directly in PyTorch. This layer enables end-to-end training without relying on external libraries. Given an input **X** ∈ ℝ^*N* ×*C*×*T*^, where *N* is the batch size, the layer computes channel-wise temporal convolutions across multiple frequency bands.

We define *S* learnable wavelet *scales* 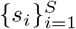, which correspond to different frequency bands and are initialized on a log scale between fixed minimum and maximum frequencies. These scales are learned during training via a parameter vector log *s*_*i*_ ∈ ℝ^*S*^. For each scale *s*_*i*_, a Morlet wavelet *ψ*_*i*_(*t*) is defined as:

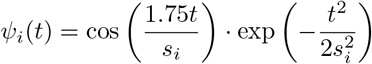

Each wavelet *ψ*_*i*_ is convolved with the input sequence independently per channel using grouped 1D convolution. A learnable weight matrix **W** ∈ ℝ^*C*×*S*^ is applied to modulate the contribution of each frequency scale per channel. The output of the wavelet layer is a tensor of shape ℝ^*N* ×*C*×*S*×*T*^, where each scale dimension captures a different band-limited representation of the input signal.

This design enables the network to learn data-driven frequency decompositions with explicit interpretability in the spectral domain, providing fine-grained insights into the dynamics of input signals across multiple frequency bands. Compared to a CNN branch as an alternative frequency-aware approach, this method offers higher interpretability due to its direct frequency associations. However, it incurs greater computational costs owing to the complexity of the transform computations.

#### 3) Encoder - Transformer Branch

In parallel, the transformer encoder branch processes the same input sequence to capture temporal dependencies and cross-feature interactions. Since self-attention is inherently permutation-invariant, we first use positional encoding to inform the model of the sequential order. We adopt absolute positional encoding for its simplicity and effectiveness. Specifically, we apply a learnable positional embedding layer before the transformer, where each position in the sequence is assigned a unique trainable vector. This vector is added to the corresponding input feature, enabling the transformer to learn temporal structure during training. Formally, for each time step *t*, the positional vector **p**_*t*_ is added to the input token **e**_*t*_, i.e., **e**_*t*_ := **e**_*t*_ + **p**_*t*_, thereby embedding absolute temporal information into the input representation.

The transformer encoder is composed of two stacked layers of multi-head self-attention (MHSA) [18] followed by positional feed-forward networks. On each layer *ℓ* (*ℓ* = 1, 2), given input token representations 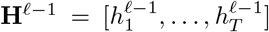 (with **H**^0^ = **E** after adding positional encodings), the self-attention computes for each head *m* = 1, 2:

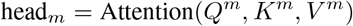

where 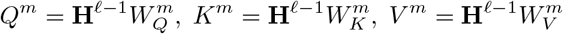

Here, 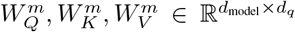 are learned projection matrices for queries, keys, and values, with *d*_*q*_ = *d*_model_*/H*.

The scaled dot-product attention for each head is:

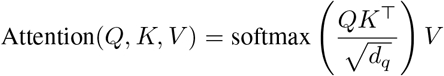

The *H* heads’outputs are then fit into temproal average pooling. This operation reduce the sequence length from *T* time steps to a single global representation. Given the transformer output 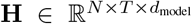 where *d*_model_ = 1378, we apply adaptive average pooling over the temporal dimension:

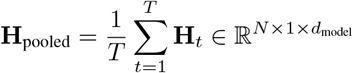

This results in a compact summary representation **H**_pooled_, which preserves the feature dimensionality while collapsing the temporal axis. It serves as a global context vector for subsequent decoding and classification.

#### 4) Encoder - Linear Branch

Given the high dimensionality of the feature space, we aim to mitigate the risk of overfitting by introducing a simple yet complementary representation. While the previously discussed frequency-aware branches already capture rich temporal and spectral dynamics, we additionally incorporate a static representation of the dFNC by applying a temporal pooling operation. This results in a static functional connectivity (sFNC) vector, which serves as a full-band summary of connectivity patterns. As a complement to the wavelet branch, this static pathway captures global structure without focusing on temporal transitions. The resulting pooled features are concatenated with the outputs from other branches and jointly passed to the decoder and classifier.

#### 5) Decoder - Reconstruction and classification

After encoding the input into a latent representation through different combinations of “expert” branches, we employ a decoder to reconstruct the original input sequence. The decoder consists of a fully connected layer that maps the latent vector back to the original input dimensions, along with a classifier for downstream prediction.

Let 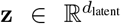 denote the latent vector obtained from the encoder. The reconstruction of the input sequence 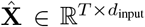 is computed as:

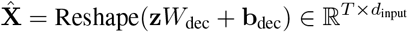

where 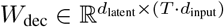 and 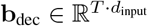 are learnable parameters, and Reshape(·) reshapes the vector into a sequence of time steps.

For classification, two separate linear projections are applied to the latent vector. The first fully connected layer reduces the dimensionality to further condense the feature representation, and the second layer produces the probability score using a sigmoid activation function.

### D. Objective Functions

For this task, we define multiple objective functions that are jointly optimized using a curriculum-based loss weighting scheduler to ensure the model is robust, sparse, and generalizable. The total objective consists of four components: reconstruction loss (ℒ_rc_), autocorrelation loss (ℒ_ac_), classification loss (ℒ_cls_), and sparsity loss (ℒ_sp_).

The reconstruction and autocorrelation losses ensure that the learned latent space faithfully represents the original input. While the reconstruction loss, implemented as mean squared error (MSE), is standard, the autocorrelation loss serves as an additional constraint to preserve the serial dependencies in the temporal structure of the reconstructed signal by enforcing similarity between time series at different time lags.

The classification loss corresponds to the primary task, where the model is trained to distinguish between two classes. We use binary cross-entropy (BCE) for this purpose.

The sparsity loss is based on the *ℓ*_1_ norm of the latent representation produced by the encoders. It encourages sparsity in the latent space, effectively performing implicit feature selection by promoting compact and informative latent encodings.

The four individual objectives are defined as:

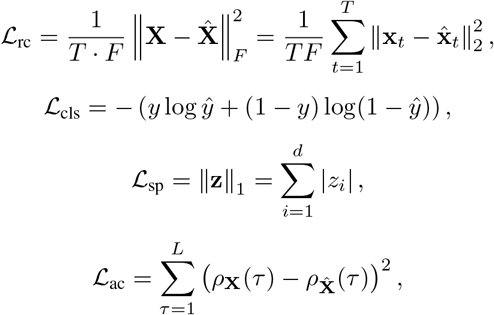

where **X** ∈ ℝ^*T* ×*F*^ is the input sequence, 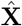 is the reconstructed sequence, **x**_*t*_ and 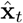 denote feature vectors at time 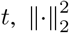 denotes the squared *ℓ*_2_ norm, **z** is the latent vector of dimension *d*, and *ρ*_**X**_(*τ*) and 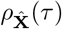 represent the autocor-relation at lag *τ* for the input and reconstructed sequences, respectively.

The total loss is a weighted sum of all components:

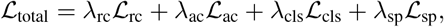

where *λ*_rc_, *λ*_ac_, *λ*_cls_, *λ*_sp_ are hyperparameters that balance the contributions of each loss term during training.

### E. Frequency-wise Model-Statistics Alignment Analysis

To gain insight into the discriminative features leveraged by the model, we visualized gradient-based saliency maps from the CNN + transformer configuration. These maps indicate how much each input feature contributes (positively or negatively) to the model’s prediction. The goal of this interpretation is to reflect group differences in actual activation levels, while also considering the model’s saliency across different frequency bands. We analyze the alignment between statistical group differences and model-learned saliency to support this objective.

Frequency-aware analysis across multiple temporal receptive fields corresponding to different convolutional kernel sizes was conducted. As described in Methods II-C.1, we used same four kernel sizes: {5, 9, 27, 49}, representing varying temporal contexts in the dFNC sequence. Each kernel size *k* was mapped to an approximate center frequency *f*_*k*_ using the formula:

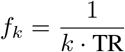

where TR is the repetition time (2.2 seconds), yielding frequency bands ranging from 0.009 to 0.091 Hz.

#### 1) Band-wise Group Difference

We computed frequency-specific group differences using dFNC data of shape [*N, C, T*]. The four kernel sizes ({5, 9, 27, 49}) defined temporal receptive fields approximating distinct frequency bands. For each kernel, we averaged dFNC over valid timepoints, resulting in a band-averaged tensor of shape [*N, B, C*], with *B* = 4 bands. To assess group-level differences between AD and CN subjects, we applied independent two-sample Welch’s t-tests [19] to each connection and frequency band. Positive t-values indicate stronger connectivity in the AD group.

#### 2) Alignment with Model Saliency

We extracted gradient-based saliency maps [20] from the trained CNN-transformer model. For each frequency band, we compared the directionality of the t-values with the corresponding saliency values. To further assess group-level differences in saliency, we also applied independent two-sample Welch’s t-tests with false discovery rate (FDR) correction (*α* = 0.01).

We defined per-band alignment using the element-wise product between the t-value matrix **T** ∈ ℝ^*B*×*C*^ and the saliency matrix **S** ∈ ℝ^*B*×*C*^ :

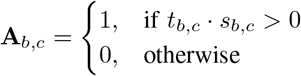

The resulting binary matrix **A** ∈ **{**0, 1}^*B*×*C*^ captures frequency-specific alignment between group-level statistical differences and model attributions. A value of **A**_*b,c*_ = 1 indicates that the model’s saliency direction is consistent with the observed group-level effect (i.e., stronger in AD or CN).

To interpret frequency-aware, model-consistent biomarkers, we masked the saliency values using this alignment matrix. The resulting masked saliency maps highlight connections that both differ statistically between groups and are relied upon by the model for prediction.

## III. Experiments

As described in the data section, we ensured that only one scan per subject was included in the dataset to prevent data leakage, which could otherwise compromise training and result reporting. We employed a 5-fold cross-validation scheme on 80% of the data for model training and validation. Each of the five trained models was then evaluated on the remaining 20% hold-out test set. To construct the hold-out set, we used stratified sampling to maintain consistent group proportions between the training and testing sets. We report model performance using the area under the receiver operating characteristic curve (ROC-AUC), which is especially appropriate given the class imbalance in the dataset. ROC-AUC evaluates the model’s ability to distinguish between classes without being overly sensitive to the specific threshold used for decision making. For fair comparison across different model configurations, we fixed the random seed during all experiments to ensure reproducibility and consistent data splits.

### A. Curriculum-based Loss Weighting

Rather than keeping fixed weights for these loss components, we employ a scheduled weighting scheme that evolves over the course of training. Formally, hyperparameters weight each loss components are non-negative weighting coefficients that change according to a predefined schedule. In the initial first 200 epochs of training, we emphasize the unsupervised objectives. We set *λ*_rc_ and *λ*_ac_ to high values so that the model prioritizes to optimize the model to efficient learn their intrinsic temporal feautres and make the reconstruction fidelity to original inputs. During this phase, the classification term and sparsity term are deliberately under-weighted (with very small *λ*_cls_ and *λ*_sp_) to prevent the model from focusing too early on the possibly noisy labels or overly sparse representations. This strategy encourages the network to learn a robust latent representation of the dFNC data, capturing meaningful neural connectivity patterns and signal dynamics without heavy pressure to immediately separate classes. After epoch 200, the training enters the second phase where the focus gradually shifts toward the supervised classification objective and the sparsity regularization. The loss weights are changed following a fixed schedule: *λ*_rc_ and *λ*_ac_ are gradually linerly decreased, while *λ*_cls_ and *λ*_sp_ are linearly increased. By epoch 200 + *M* (towards the end of training), the classification loss and sparsity penalty dominate the total loss. In our experiments, we set *M* (the transition duration) based on validation performance for early stopping. Curriculum-based loss weighting, defined a priori, progressively emphasizes different learning objectives to support staged model training.

## IV. Results and Interpretation

### A. Alzheimer’s Disease Classification Performance

We evaluate the effectiveness of FINE for AD classification using dFNC features across different combinations of expert branches. To assess the contribution of each architectural component, we conduct an ablation study where individual branches (e.g., linear, transformer, wavelet, CNN) and selected combinations are enabled. We report the area under the AUC with mean and standard deviation over five folds.

Figure 3 summarizes the classification performance of each model configuration. Among individual expert branches, the wavelet module achieves the highest standalone performance with a mean ROC-AUC of 0.749 ± 0.033, demonstrating its ability to extract informative frequency-specific biomarkers. The transformer branch also performs competitively (0.730 ± 0.021), indicating that modeling temporal dependencies is beneficial. In contrast, the linear branch shows lower discriminative power (0.698 ± 0.047), consistent with its limited capacity for capturing complex dynamics.

**Fig. 3:**
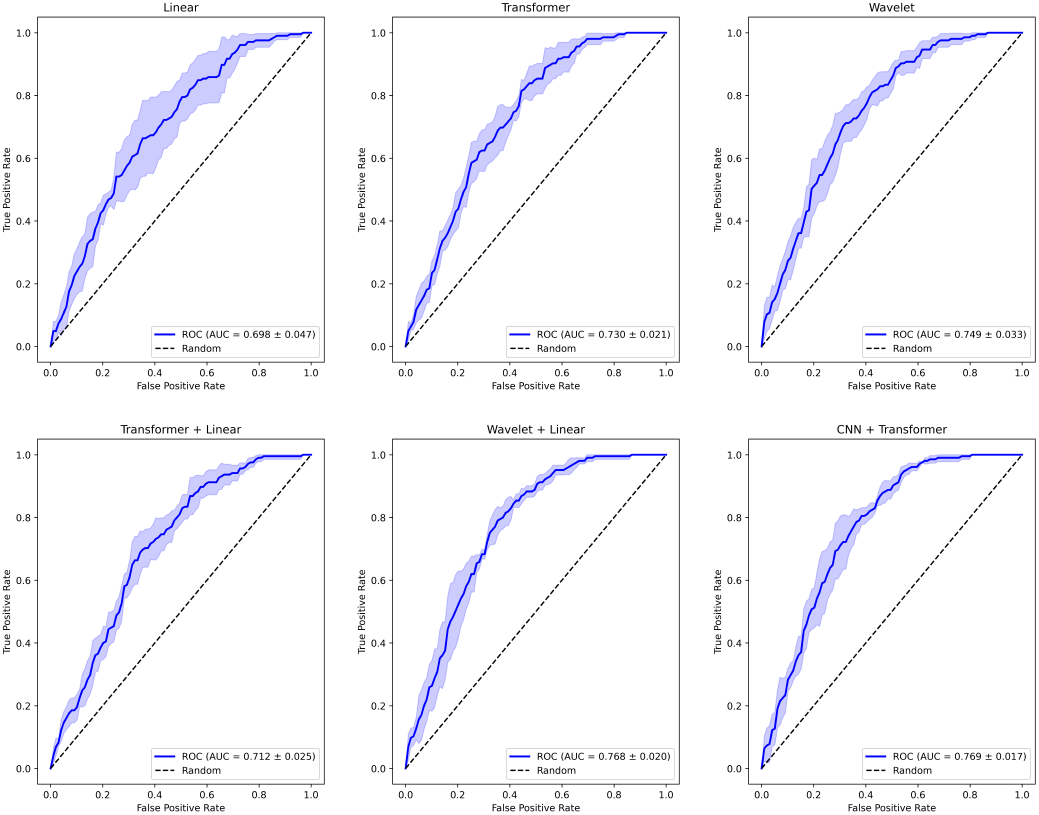
Ablation study comparing ROC curves for different combinations of expert branches in FINE. Each plot shows the mean ROC-AUC and standard deviation across 5-fold cross-validation. Frequency-aware modules (wavelet, CNN) and temporal encoders (transformer) contribute significantly to classification performance. The best performance is achieved by combining CNN and transformer branches.

Combining structural and frequency-aware branches further enhances performance. The combination of CNN and transformer achieves the highest AUC among all tested configurations (0.769 ± 0.017), suggesting that these branches capture complementary information across temporal and spectral domains. Similarly, the wavelet-linear configuration yields strong performance (0.768 ± 0.020), highlighting the benefit of integrating frequency features with wavelet transform with structural context. Interestingly, the transformer-linear pairing (0.712 ± 0.025) provides a modest gain over linear alone, but underperforms compared to frequency-aware combinations. This emphasizes the critical role of frequency representation in AD classification. These results validate the core design principle of FINE: incorporating modular, frequency-sensitive and structural-aware branches leads to improved classification performance.

### B. Multi-objective Loss Performance

Table III presents the quantitative comparison of six models trained with the curriculum-based loss weighting schedule. The CNN-transformer architecture achieved the best overall performance across all three loss metrics, recording the lowest reconstruction loss (0.0757), the lowest autocorrelation loss (0.0082), and the lowest sparsity loss (0.0956). This demonstrates the strength of combining convolutional modules, which capture local temporal-frequency patterns, with transformers, which excel at modeling global dependencies.

**TABLE III:** Comparison of the performance of all six models on three loss functions: reconstruction loss, autocorrelation loss, and sparsity loss averaged across folds under the Curriculum-based Loss Weighting strategy.

| Model | Reconstruction Loss | Autocorrelation Loss | Sparsity Loss |
| --- | --- | --- | --- |
| Linear | 0.0887 ± 0.0007 | 0.0105 ± 0.0003 | 0.1719 ± 0.0000 |
| Transformer | 0.0988 ± 0.0021 | 0.0124 ± 0.0004 | 0.1275 ± 0.0067 |
| Wavelet | 0.0902 ± 0.0017 | 0.0104 ± 0.0003 | 0.7796 ± 0.0084 |
| Linear + Transformer | 0.0951 ± 0.0092 | 0.0109 ± 0.0009 | 0.1543 ± 0.0079 |
| Linear + Wavelet | 0.0895 ± 0.0010 | 0.0109 ± 0.0005 | 0.6854 ± 0.0036 |
| CNN + Transformer | **0.0757 ± 0.0022** | **0.0082 ± 0.0005** | **0.0956 ± 0.0201** |

Interestingly, wavelet-based models (both the standalone and combined with linear components) consistently performed well on the autocorrelation loss, suggesting their inherent ability to preserve temporal structure through multiscale frequency decomposition. The wavelet-involved model exhibited the highest sparsity loss, which arises from the nature of the wavelet transform: it uses multiple scaled and shifted filters to cover the entire signal, yielding an overcomplete representation that does not form a minimal compact basis.

The linear baseline achieved strong reconstruction performance, but underperformed in sparsity and classification-related behavior, likely due to its limited modeling capacity. transformer-only and linear-transformer models performed reasonably across all metrics but did not outperform the frequency-aware variant, indicating the importance of localized frequency-related structure modeling in addition to global context.

It is worth noting that the reconstruction and autocorrelation losses were prioritized during the early training phase. This design inherently encourages all models to initially focus on unsupervised objectives. As a result, these losses typically reached their best levels before epoch 200 and could increase slightly afterward as the model gradually shifts attention toward classification and sparsity. This trade-off is a direct consequence of shifting loss weighting and reflects the intended balance between learning robust representations and ultimately optimizing for supervised discrimination. Overall, these results validate the effectiveness of different combination of expert branches’ across different learning objective.

### C. Saliency-based Frequency-wise Interpretability

We visualized frequency-specific group differences supported by both statistical tests and model derived saliency, highlighting functional connections that are not only significantly different between groups but also contribute strongly to the model’s predictions. As shown in Figure 4, distinct and structured patterns emerge across the four frequency bands.

**Fig. 4:**
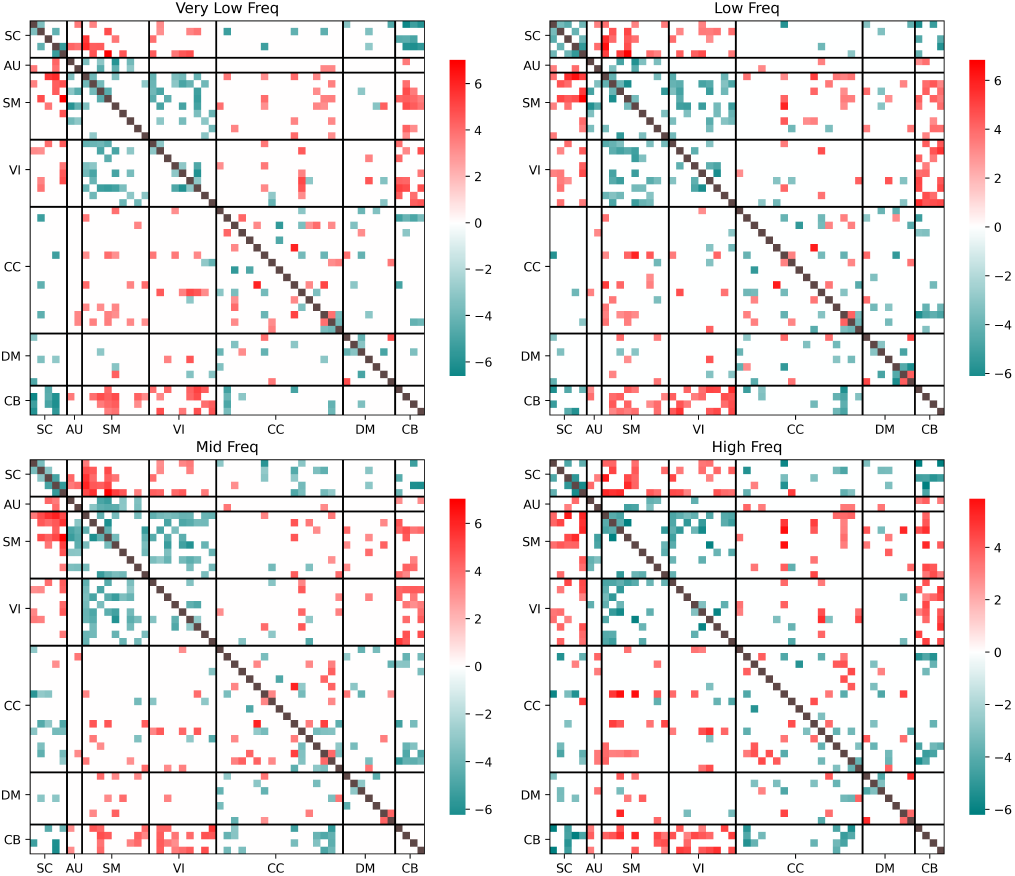
Model-aligned t-values across four frequency bands. Each matrix shows the connection-wise t-values masked by alignment with the model’s saliency (i.e., only showing values where the model and group statistics agree in direction). Positive values (red) indicate stronger connectivity in AD; negative values (blue) indicate stronger connectivity in CN.

The aligned t-value maps across frequencies reveal several consistent and frequency-specific patterns that correspond well with known AD-related brain alterations. Across all bands, consistent trends are observed: reduced connectivity in SC–SC, SC–CB, SM–SM, SM–AU, SM–VI, and CB–CC pathways in the AD group. These patterns align well with prior findings, [21] reported decreased SC–CB connectivity in AD, while [8] highlighted reduced SM–VI activity. Conversely, the observed increase in VI–CB connectivity is also consistent with [21], further supporting a potential compensatory mechanism. Those findings highlighting their potential as functional connectivity biomarkers

In our results, the low and mid-frequency bands exhibit more significant group differences compared to the very low and high-frequency bands. This trend is consistent with findings from multiple ALFF-based studies, which report greater sensitivity to AD related changes within the 0.01–0.1 Hz range, typically associated with physiologically meaningful slow neural oscillations.

At low frequencies, prominent differences are concentrated in the thalamus within the SC domain, which plays a key role in sensorimotor integration and is often implicated in early AD progression. This observation aligns with [22], which reported altered thalamic activity in AD using ALFF metrics within the 0.01–0.1 Hz range.

At mid and high frequencies, reduced activity is more pronounced within SM–SM connections, particularly involving the postcentral gyrus, a region responsible for somatosensory processing. This is consistent with findings in [23], further supporting frequency-dependent degeneration in motor-related networks. Similarly, CC–SC connections also show greater disruption in the mid-frequency range compared to lower frequencies, suggesting a decline in executive subcortical integration with disease progression. In contrast, activity within the middle temporal gyrus, associated with language and semantic memory, becomes more diffuse and less frequency-localized, indicating a possible breakdown in frequency-specific functional specialization in higher-order cognitive regions.

### D. Interpreting Frequency Prioritization in Wavelet Models

Unlike fixed CNN kernels, the learnable wavelet layer adjusts its scale parameters during training, allowing it to tune frequency responses based on the data. Analysis of the trained models, including both the wavelet–linear combination and the standalone wavelet model, shows that the learned scales consistently emphasize frequencies around 0.03 Hz and 0.1 Hz. These frequencies correspond to the low and mid-frequency bands and are well aligned with prior studies identifying them as informative for detecting Alzheimer’s-related alterations in brain connectivity [21], [22].

Interestingly, the learned scales also include frequencies that fall outside the conventional resting-state fMRI range (e.g., below 0.01 Hz). This behavior stems from the flexibility of the wavelet design, which allows the model to adapt its filters freely based on the interaction of multiple loss objectives. In particular, the sparsity loss encourages the network to form compact representations, often driving the model to select a sparse subset of informative frequencies. Frequencies like 0.03 Hz and 0.1 Hz, which lie at the center of low and mid-frequency ranges, may serve as proxies for neighboring bands due to their representative signal characteristics. As a result, redundant frequencies in close proximity may be suppressed to meet the sparsity objective, consistent with the frequency redundancy we observed in earlier saliency-based interpretations.

Some learned frequencies fall outside the typical 0.01 to 0.15 Hz range used in resting-state fMRI analysis. This may reflect the flexibility of the wavelet layer and interactions with sparsity constraints, though such low frequencies can also be influenced by scanner drift and are generally considered physiologically less meaningful [24].

## V. Discussion and Conclusion

In this work, we introduced FINE, a frequency-aware, interpretable deep learning model designed to address key limitations in the analysis of dFNC for AD classification. Through its modular architecture, FINE captures complementary spectral and temporal features while supporting rigorous interpretability at the frequency-band level.

Our results reveal several important findings. First, frequency-aware modeling is crucial for achieving high classification performance. Both the wavelet and CNN branches outperformed purely temporal or static baselines, underscoring the clinical relevance of frequency-specific features in AD-related brain dynamics. Notably, the combination of CNN and transformer branches achieved the best overall ROC-AUC, highlighting that local frequency patterns and long-range temporal dependencies are synergistic for AD prediction.

Second, our frequency-wise model-statistics alignment analysis provides compelling evidence that FINE learns potential biologically meaningful representations. The identified frequency-specific connectivity disruptions, including reduced connections such as SC–CB, SM–VI, SM-SM and elevated VI–CB interactions, align well with prior findings in the literature and with known pathophysiological mechanisms in AD. Importantly, we observed that these disruptions are not uniform across the spectral domain: low and mid-frequency bands exhibited stronger and more consistent group differences, consistent with prior ALFF studies and the known role of slow neural oscillations in cognition and disease progression.

From an interpretability standpoint, our saliency-based masking approach provides a principled way to reconcile data-driven deep learning attributions with classical statistical group analyses. By focusing on model-aligned features, we ensure that interpretations reflect both statistical group differences and the model’s decision process. This approach mitigates the risk of highlighting spurious or overfit features, an often underappreciated challenge in XAI for neuroimaging.

There are several broader implications of this work. First, our findings emphasize the value of moving beyond full-band dFNC analysis. Conventional approaches that collapse across the frequency domain may obscure critical biomarkers that are frequency-selective. Second, FINE provides a flexible framework for investigating other brain disorders where frequency-specific alterations are hypothesized (e.g., schizophrenia, depression). Its modular design supports the incorporation of additional expert branches, such as graph-based encoders or cross-modal fusion layers, enabling future extension to multimodal neuroimaging.

Moreover, the explicit modeling of temporal and spectral dynamics in an interpretable manner bridges an important gap between deep learning and neuroscientific insight. While prior models have achieved strong classification performance, few have provided actionable hypotheses regarding how and where disease impacts brain dynamics across scales. FINE contributes toward this goal by offering both predictive power and mechanistic transparency.

Several limitations call for consideration. First, although we validated FINE on a large AD dataset, future work should evaluate its generalizability to independent cohorts and longitudinal disease progression. Second, while our CNN-based frequency modeling is effective, its mapping to precise physiological frequency bands remains approximate and limit. Future research could incorporate more principled approaches for defining frequency bands. Finally, while we focused on group differences that align with model-derived saliency, there are also connections where the model assigns importance but the direction of effect contradicts the group-level t-values. This discrepancy may reflect model overfitting, but it could also arise from complex nonlinear interactions not captured by standard statistical tests. These patterns warrant further investigation to enhance interpretability.

In conclusion, FINE demonstrates that frequency-aware, interpretable deep learning can advance both disease classification and neuroscientific understanding of dynamic brain connectivity in AD. By bridging spectral, temporal, and spatial modeling in a transparent architecture, FINE offers a promising foundation for future work in explainable neuroimaging and precision diagnostics.

## Acknowledgment

This research supported by NSF 2112455 and NIH R01AG073949.

